# A sex-stratified genome-wide association study of tuberculosis using a multi-ethnic genotyping array

**DOI:** 10.1101/405571

**Authors:** Haiko Schurz, Craig J Kinnear, Chris Gignoux, Genevieve Wojcik, Paul D van Helden, Gerard Tromp, Brenna Henn, Eileen G Hoal, Marlo Möller

## Abstract

Tuberculosis (TB), caused by Mycobacterium tuberculosis, is a complex disease with a known human genetic component. Males seem to be more affected than females and in most countries the TB notification rate is twice as high in males as in females. While socio-economic status, behaviour and sex hormones influence the male bias they do not fully account for it. Males have only one copy of the X chromosome, while diploid females are subject to X chromosome inactivation. In addition, the X chromosome codes for many immune-related genes, supporting the hypothesis that X-linked genes could contribute to TB susceptibility in a sex-biased manner. We report the first TB susceptibility genome-wide association study (GWAS) with a specific focus on sex-stratified autosomal analysis and the X chromosome. Individuals from an admixed South African population were genotyped using the Illumina Multi Ethnic Genotyping Array, specifically designed as a suitable platform for diverse and admixed populations. Association testing was done on the autosome and X chromosome in a sex stratified and combined manner. SNP association testing was not statistically significant using a stringent cut-off for significance but revealed likely candidate genes that warrant further investigation. A genome wide interaction analysis detected 16 significant interactions. Finally, the results highlight the importance of sex-stratified analysis as strong sex-specific effects were identified on both the autosome and X chromosome.

## Introduction

Tuberculosis (TB) caused by *Mycobacterium tuberculosis* (*M. tuberculosis*) is a global health epidemic and the leading cause of death due to a single infectious agent (1). In 2016 1.3 million TB deaths were reported in HIV negative individuals and an additional 374000 deaths related to TB/HIV co-infection were recorded. The majority of these deaths occurred in southeast Asian and African countries (1). TB is a complex disease, influenced by environmental and behavioural factors such as socio-economic status and smoking, as well as definite human genetic components. The contribution of the host genes to disease has been highlighted by numerous investigations, including animal (2), twin (3–5), linkage (6,7) and candidate gene association studies (8). More recently genome-wide association studies (GWAS) in diverse populations have been done (9–18).

Interestingly another influential factor in TB disease development is an individual’s biological sex, which has been largely ignored in past TB studies and was usually only used as a covariate for adjusting association testing statistics. In 2016, males comprised 65% of the 10.4 million recorded TB cases, indicating that the TB notification rate is nearly twice as high in males as in females (WHO, 2017). While socio-economic and behavioural factors do influence this ratio, it does not fully explain the observed sex-bias (19). Another factor that influences sex-bias is the effect that sex hormones (estrogen and testosterone) have on the immune system. Estrogen is an immune activator, upregulating pro-inflammatory cytokines (TNFα), while testosterone is an immune suppressor, upregulating anti-inflammatory cytokines (IL-10) (20). This could explain why men are more susceptible to infectious diseases compared to females (19). However, as sex-based differences in immune responses differ even between pre-pubertal boys and girls, as well as between post-menopausal women and elderly men, it shows that sex hormones do not fully explain the sex-bias (21). Thus, it has been proposed that the X chromosome and X-linked genes directly contribute to the observed sex-bias.

There are approximately 1500 genes on the X chromosome, many of which are involved in the adaptive or innate immune system (22). Since females have two X chromosomes, one requires silencing in order to equalise dosage of gene expression to that of men who only have one X chromosome. This silencing occurs randomly in each cell, making females functional mosaics for X linked genes and giving them a major immunological advantage over males (19). As males are haploid for X-linked genes any damaging polymorphisms or mutations on the X chromosome will have a more pronounced immunological effect in males than in mosaic females, thereby influencing the sex-bias (23).

To date, eleven GWAS investigating susceptibility to clinical TB have been published (9–18,24). There has not been significant overlap between the 11 published TB GWAS, but it seems that replication is more likely when populations with similar genetic backgrounds are compared: the *WT1* locus was associated with disease in populations from West and South Africa (9,17). Critically, genotyping microarrays that did not fully accommodate African genetic diversity were used in these studies (9–11,17,18). It is therefore possible that unique African-specific susceptibility variants were not tagged by these initial arrays, since linkage disequilibrium (LD) blocks are shorter in African populations (25). Moreover, none of the GWAS included or examined the X chromosome or sex-stratified analysis of the autosomes as was done in an asthma cohort (26). Genetic differences between asthmatic males and females were identified on the autosome, with certain alleles having opposite effects between the sexes. Independent confirmation of the involvement of the X chromosome in TB susceptibility is the association of X-linked *TLR8* susceptibility variants with active TB. Davila *et al.* (27) investigated 4 *TLR8* variants (rs3761624, rs3788935, rs3674879, rs3764880) in an Indonesian cohort and showed that all variants conferred susceptibility to TB in males but not females. The results for males were validated in male Russian individuals (27). These results were validated for rs3764880 in Turkish children, but no significant association was found for rs3764879 (28). Hashemi-Shahri *et al.* (29) found no significant *TLR8* associations in an Iranian population, while rs3764880 was significantly associated with TB susceptibility in both males and females in a Pakistani cohort (30). In South African Coloured (SAC) individuals rs3764879 and rs3764880 were significantly associated in both males and females, while rs3761624 was only significantly associated in females (31). Interestingly, in this cohort opposite effects were consistently found between the sexes for the same allele in all investigated *TLR8* variants (31), echoing the asthma findings of Mersha et al (26). Finally, in a Chinese cohort rs3764879 was significantly associated with TB disease in males but not females.

We report the first TB susceptibility GWAS with a specific focus on sex-stratified autosomal analysis and the X chromosome to elucidate the male sex-bias. This is also the first GWAS in an admixed South African population that uses an array (Illumina Multi Ethnic Genotyping Array) specifically designed to detect variants in the 5 most commonly studied populations, making it the most suitable platform for diverse and admixed populations.

## Materials and methods

### Study population

Study participants were recruited from two suburbs in the Cape Town metropole of the Western Cape. These suburbs were chosen for its high TB incidence and low HIV prevalence (2%) at the time of sampling (1995-2005) (32). Approximately 98% of the residence in these suburbs self-identify as SAC and have the same socio-economic status, which reduces confounding bias in the association testing (9). The cohort consists of 420 pulmonary TB (pTB) cases, bacteriologically confirmed to be culture and/or smear positive and 419 healthy controls from the same suburbs. Approximately 80% of individuals over the age of 15 years from these suburbs have a positive tuberculin skin test (TST), indicating exposure to *M. tuberculosis* (33). All study participants were over 18 years of age and HIV negative.

Approval was obtained from the Health Research Ethics Committee of Stellenbosch University (project registration number S17/01/013 and 95/072) before participant recruitment. Written informed consent was obtained from all study participants prior to blood collection. DNA was extracted from the blood samples using the Nucleon BACC Genomic DNA extraction kit (Illumina, Buckinghamshire, UK). DNA concentration and purity was checked using the NanoDrop^®^ ND-1000 Spectrophotometer and NanoDrop^®^ v3.0.1 software (Inqaba Biotechnology, Pretoria, SA). The study adhered to the ethical guidelines as set out in the “Declaration of Helsinki, 2013 (34).

### Genotyping

Genotyping was done using the Illumina Multi-ethnic genotyping array (MEGA) (Illumina, Miami, USA) which has content from various ethnicities making it highly suitable for diverse and admixed populations. Genome studio v2.04 (Illumina, Miami, USA) was used for SNP calling to calculate intensity scores and to call common variants (MAF >= 5%), followed by analysis with zCall to recall rare genotypes (MAF <5%) (35).

### Genotyping quality control

Quality control (QC) of the genotyping data was done using the XWAS version 2.0 software and QC pipeline to filter out low quality samples and SNPs (36,37). Data were screened for sex concordance, relatedness (up to third degree of relatedness) and population stratification (as determined by principal component analysis). Genotypes for males and females were filtered separately in order to maintain inherent differences between the sexes. SNPs were removed from the analysis if missingness correlated with phenotype (threshold = 0.01) as well as individual and SNP missingness (greater than 10%), minor allele frequency (less than 1%) and Hardy–Weinberg equilibrium (HWE) in controls (threshold = 0.01). Filtering continued iteratively until no additional variants or individuals were removed. Overlapping markers between the sexes were merged into a single dataset. X chromosome genotypes were extracted and variants were removed if the MAF or missingness was significantly different between the sexes (threshold = 0.01). A flow diagram explaining quality control steps and association testing of the data is shown in Figure s1.

### Admixture

The SAC population is a 5-way admixed population with ancestral contributions from Bantu-speaking African populations, KhoeSan, Europeans and South and East Asians. To avoid confounding during association testing the ancestral components are included as covariates (38). Admixture was estimated for the autosome (chromosome 1-22) and the X chromosome separately using the software ADMIXTURE (v1.3) (39) and reference genotyping data for 5 ancestral populations. The reference populations used to infer ancestry were European (CEU) and South Asian (Gujarati Indians in Houston, Texas and Pathan of Punjab) extracted from the 1000 Genomes Phase 3 data (40), East Asian (Han Chinese in Beijing, China), African (Luhya in Webuye, Kenya, Bantu-speaking African, Yoruba from Nigeria) and San (Nama/Khomani) (41,42). SAC samples were divided into running groups in order to keep the number of individuals per reference population and admixed population consistent. Each running group was analysed five times at different random seed values. The results for each individual were averaged across the five runs in order to obtain the most accurate ancestry estimations (43). Four ancestral components (African, San, European and South Asian(44)) were included as covariates in the logistic regression association testing with the smallest component (East Asian) excluded in order to avoid complete separation of the data.

### Association analysis

#### SNP based association analysis

Autosomal TB association testing was done with sex-stratified and combined datasets using the additive model in PLINK (version 1.7^1^) (45) in order to detect sex-based differences. TB association testing for the X chromosome were done separately in males and females using XWAS (version 2) and the results were combined using Stouffers method in order to obtain a combined association statistic (36,37). A sex-differentiated test was conducted for the X chromosome using the XWAS software to test for significant differences in genetic effects between males and females. X chromosome inactivation states were also included in the association testing as covariates using a method developed by Wang *et al.* (46). Ancestry, sex and age were included in analyses as covariates where applicable. Multiple testing correction was done using the SimpleM method (47), which adjusts the significance threshold based on the number of SNPs that explains 95% of the variance in the study cohort. This method is less conservative than Bonferroni correction and is a close approximation of permutation results in a fraction of the time. For the autosome the genome-wide significance threshold was set to 5.0e^-8^ (48).

### Gene based association analysis

Gene-based association testing groups SNPs together and thus decreases the multiple testing burden and increase power to detect an association. Gene-based association testing was done using the XWAS v2 scripts, which were implemented using the Python^2^ (version 2.7.10) and R programming environment (version 3.2.4, (49)) and R packages corpcor and mvtnorm. Reference files for the known canonical genes on the X chromosome for human genome build 37 were included in the XWAS v2 software package and used to group variants and p-values by gene (36,37). Bonferroni correction was used to adjust for multiple testing to minimise false positive associations.

### Interaction analysis

Genome-wide SNP interaction analysis was done using CASSI^3^ (v2.51). A joint effects model was implemented for a rapid overview of interactions of all variants across the genome (autosome and X chromosome). Variants from significant interactions were reanalysed using a logistic regression approach with covariate correction, which would not be feasible for a genome-wide interaction analysis as it would be too computationally intensive. Bonferroni correction was done for the number of interactions tested to avoid false positives.

## Results

### Cohort summary

In total 410 TB cases and 405 healthy controls passed the sex-stratified QC procedure. General summary statistics for the cohort, including mean and standard deviation of age and global ancestry as well as the ratio of males to females in both cases and controls are shown in Table 1. Clear differences were observed between TB cases and controls for both age and ancestry, justifying the inclusion as covariates. Ancestral distributions also differed between the sexes and between the autosome and X chromosome (Figure 1) and as a result, the autosomal or X chromosome ancestral components were included as covariates in the respective analyses.

**Table 1:** SAC sample characteristics showing case/control and sex distribution, mean and standard deviation of age and global ancestral components

| Group | Number | Female (%) | Age | San | African | European | South Asian | East Asian |
| --- | --- | --- | --- | --- | --- | --- | --- | --- |
| **TB cases** | 410 | 242 (59) | 36.32 ± 11.04 | 33.89 ± 18.83 | 29.11 ± 19.80 | 17.10 ± 16.81 | 12.95 ± 10.82 | 7.08 ± 7.22 |
| **Controls** | 405 | 223 (55) | 30.55 ± 12.91 | 33.75 ± 19.59 | 29.92 ± 20.46 | 16.12 ± 15.81 | 13.26 ± 12.38 | 7.04 ± 7.44 |

**Figure 1:**
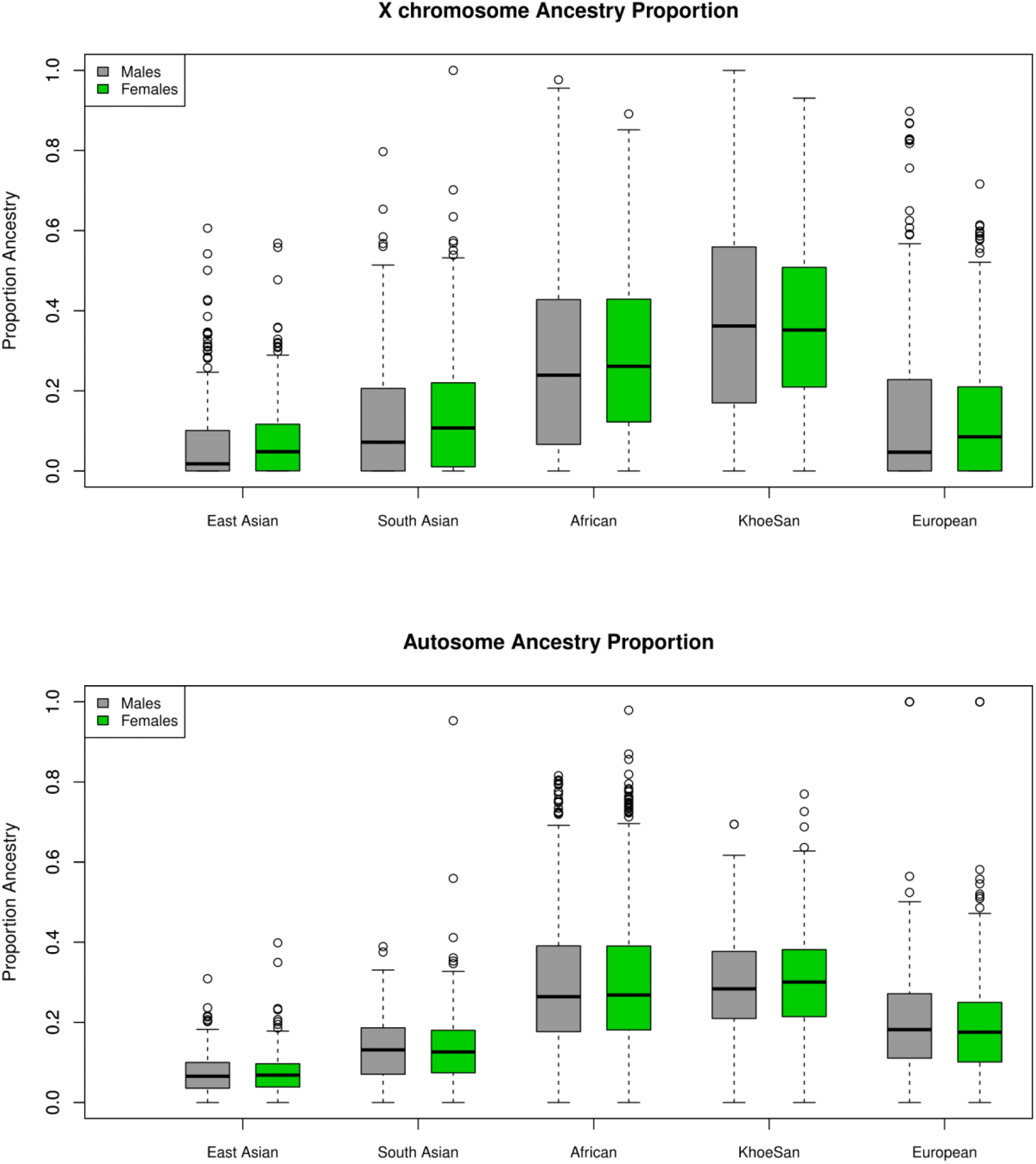
Ancestral distribution on the X chromosome and autosome for males and females.

## Association testing results

### SNP based

The top results for the autosomal association testing are shown in Table 2 and Figure S2, with the QQ-plot figindicating no constraints on the analysis or inflation of the results (Figure S2). Following multiple test correction, no significant associations were identified for the combined or sex-stratified analysis, but it is important to note that the top associations differed between the sex-stratified and combined analyses as well as for males and females (Table 2). The most significant variant for the combined autosomal association test was rs17410035 (OR= 0.4, p-value = 1.5e^-6^, table 2), located in the 3’-UTR of the *DROSHA* gene, which encodes a type 3 RNase. This RNase is involved in miRNA processing and miRNA biogenesis (50). Although little evidence exists that rs17410035 has an impact on *DROSHA* gene expression or miRNA biogenesis (which could affect gene expression) it has been associated with increased colon cancer (OR= 1.22, p-value = 0.014) (50) and cancer of the head and neck (OR= 2.28, p-value = 0.016) (51). When the rs17410035 SNP interacts with other variants (rs3792830, rs3732360) it can further increase the risk for cancer of the head and neck (51), which illustrates the importance of doing interaction analysis. For the autosomal sex-stratified analysis the most significant association in males was rs11960504 (OR = 2.8, p-value = 7.21e^-6^, table 2) located downstream of the *GRAMD2B* gene, a gene for which no information is available. The most significant SNP in the females was rs2894967 (OR = 2.17, p-value = 4.77e^-6^) located upstream of the *TENT4A* gene, a gene coding for a DNA polymerase shown to be involved in DNA repair (52). Closer inspection of the data revealed that the effects between the sexes were in the same direction for all top hits in the combined analysis, whereas all variants identified in the sex-stratified analysis had effects in opposite directions between the sexes, or one sex had no effect, indicating that even on the autosome strong sex specific effects are prominent.

**Table 2:** Top associations for the combined and sex-stratified autosomal association testing.

| Chr | SNP | A1 | Location | Gene | Group | OR | 95CI | P-value |
| --- | --- | --- | --- | --- | --- | --- | --- | --- |
| 5 | rs17410035 | T | 5'UTR | *DROSHA* | Combined | 0.404 | 0.28-0.58 | $1.50 \times 10^{-6}$ |
| 5 | rs1501847 | G | 5'UTR | *C5orf64* | Combined | 1.708 | 1.37-2.14 | $2.64 \times 10^{-6}$ |
| 7 | rs2665441 | C | 3'UTR | *ASNS* | Combined | 1.681 | 1.34-2.10 | $5.51 \times 10^{-6}$ |
| 9 | rs1662230 | G | 5'UTR | *RN7SKP120* | Combined | 2.278 | 1.58-3.27 | $8.91 \times 10^{-6}$ |
| 12 | rs199911028 | G | Intronic | *CFAP54* | Combined | 2.966 | 1.89-4.67 | $2.58 \times 10^{-6}$ |
| 15 | rs142644068 | C | Intronic | *PCSK6* | Combined | 0.132 | 0.06-0.30 | $1.56 \times 10^{-6}$ |
| 5 | rs11960504 | T | 3'UTR | *GRAMD2B* | Male | 2.801 | 1.79-4.39 | $7.21 \times 10^{-6}$ |
| 13 | rs9315991 | A | Intronic | *LINC00400* | Male | 0.394 | 0.27-0.58 | $2.03 \times 10^{-5}$ |
| 14 | rs8016621 | A | Intronic | *SALL2* | Male | 0.252 | 0.14-0.46 | $5.91 \times 10^{-6}$ |
| 5 | rs2894967 | C | 5'UTR | *TENT4A* | Female | 2.173 | 1.56-2.90 | $4.77 \times 10^{-6}$ |
| 9 | rs10819610 | T | Intronic | *NCS1* | Female | 0.514 | 0.39-0.67 | $1.55 \times 10^{-6}$ |
| 14 | rs7152005 | T | Intronic | *DPF3* | Female | 2.13 | 1.66-2.90 | $1.52 \times 10^{-6}$ |
| 21 | rs2150367 | T | Intronic | *LINC02246* | Female | 0.502 | 0.38-0.67 | $1.56 \times 10^{-6}$ |

For the X chromosome specific association testing a sex-stratified test was conducted and the results were then combined using Stouffers method, which provided a good fit between expected and observed p-values (QQ-plot Figure 3) (36,37). The simpleM method indicated that of the 20939 X-linked variants 17600 explained 95% of the variance in the data resulting in a significance threshold of 2.8e^-6^ (0.05/17600). No statistically significant associations with TB susceptibility were identified in either sex-stratified or the combined analysis (Table 3 and Figure 3). The most significant association for the X-linked combined (p-value = 2.62e^-5^) and females (OR = 1.83, p-value = 1.06e^-4^) only analysis was the same variant, rs768568, located in the *TBL1X* gene. For the males the most significant association was rs12011358 (OR = 0.37, p-value = 1.25e^-4^) located in the *MTND6P12* gene. Both of these genes have not been previously associated with TB susceptibility and *MTND6P12* is a pseudogene with unknown expression patterns or function. Variants in *TBL1X* have been shown to influence prostate cancer (53) and central hypothyroidism (54) susceptibility. *TBL1X* is a regulator of nuclear factor kappa-light-chain-enhancer of activated B cells (NF-kB) and is thus involved in the immune system which could impact TB susceptibility.

**Table 3:** Most significant X-linked associations, using Stouffers method to combine p-values.

| Chr | SNP | A1 | Location | Gene | Male |  |  | Female |  |  | P_Diff |
| --- | --- | --- | --- | --- | --- | --- | --- | --- | --- | --- | --- |
|  |  |  |  |  | OR | 95CI | P | OR | 95CI | P |  |
| X | rs145407087 | C | 3'UTR | *RNU6-974P* | 0.427 | 0.12-1.54 | 0.193 | 7.147 | 1.95-26.19 | 0.003 | 8.33e<sup>-5</sup> |
| X | rs5917743 | C | 5'UTR | *SRPX* | 0.300 | 0.05-1.89 | 0.203 | 15.04 | 1.81-124.7 | 0.012 | 2.18e<sup>-4</sup> |
| X | rs1337567 | C | 5'UTR | *DIAPH2* | 1.45 | 0.88-2.39 | 0.146 | 0.571 | 0.42-0.78 | 3.49e<sup>-4</sup> | 6.73e<sup>-4</sup> |

**Table 3:**
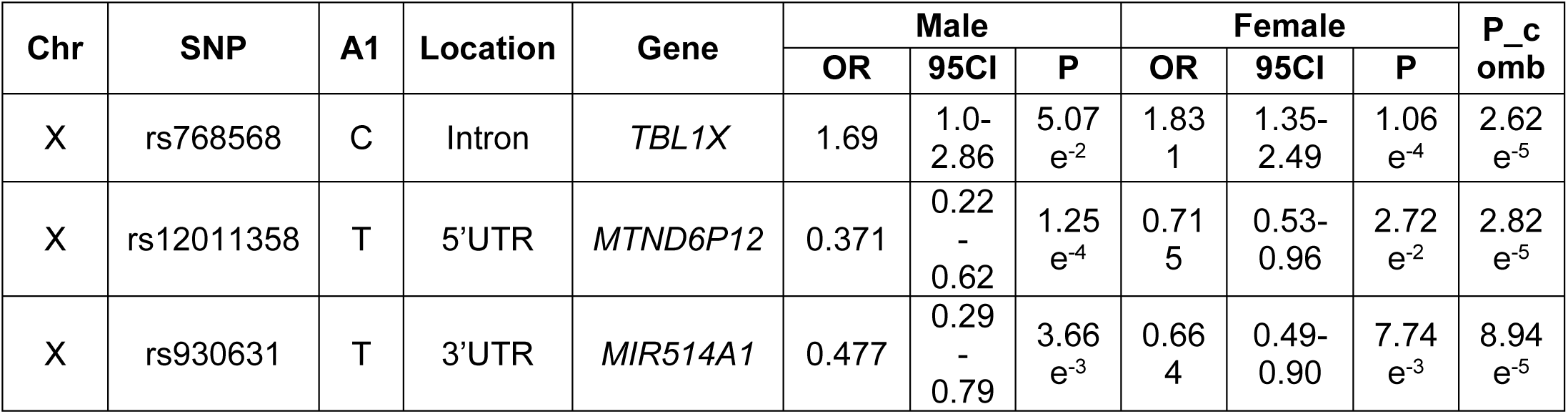
Sex-differentiation analysis.

| Chr | SNP | A1 | Location | Gene | Male |  |  | Female |  |  | P_c<br>omb |
| --- | --- | --- | --- | --- | --- | --- | --- | --- | --- | --- | --- |
|  |  |  |  |  | OR | 95CI | P | OR | 95CI | P |  |
| X | rs768568 | C | Intron | *TBL1X* | 1.69 | 1.0-2.86 | 5.07<br>e<sup>-2</sup> | 1.83<br>1 | 1.35-2.49 | 1.06<br>e<sup>-4</sup> | 2.62<br>e<sup>-5</sup> |
| X | rs12011358 | T | 5'UTR | *MTND6P12* | 0.371 | 0.22<br>-<br>0.62 | 1.25<br>e<sup>-4</sup> | 0.71<br>5 | 0.53-0.96 | 2.72<br>e<sup>-2</sup> | 2.82<br>e<sup>-5</sup> |
| X | rs930631 | T | 3'UTR | *MIR514A1* | 0.477 | 0.29<br>-<br>0.79 | 3.66<br>e<sup>-3</sup> | 0.66<br>4 | 0.49-0.90 | 7.74<br>e<sup>-3</sup> | 8.94<br>e<sup>-5</sup> |

The method of modelling X chromosome inactivation states, developed by Wang *et al.* (46), was also incorporated into the X-linked association testing, but no significant observations were observed. Although the p-values were generally lower than for the Stouffer method, the QQ-plot revealed that including estimations of X chromosome inactivation states inflated the p-values and increased the chance of type 1 errors and these results were therefore discounted (Table S1 and Figure S3).

The sex differentiation test did not result in any significant associations (Table 4), with the most significant association in a pseudogene, *RNU6-974P* (p-value = 8.33e^-5^). The second most significant association was for a variant upstream of the *SRPX* (p-value = 2.18e^-4^) gene which has previously been shown to have a tumour suppressor function in prostate carcinomas (55). Whether these variants are associated with TB susceptibility or influence sex-bias is unclear, but the vastly opposite effects between the sexes are noteworthy. When comparing the OR for the sex differentiation test it is clear that variants can have major sex specific effects again highlighting the need for sex-stratified analysis (Table 4).

**Table 4:**
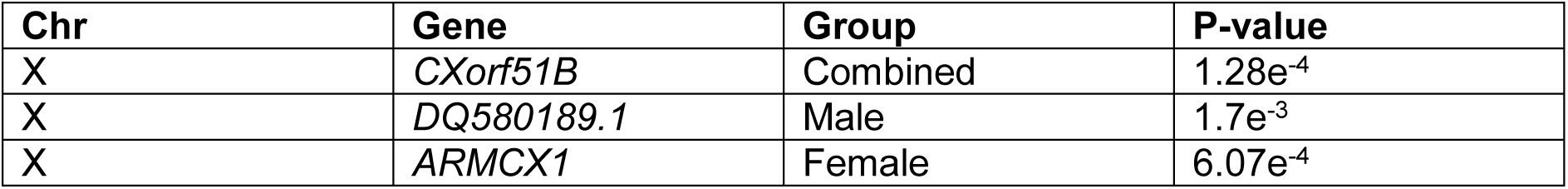
X chromosome gene-based association results.

### Gene based

The X chromosome gene-based analysis, in which 1105 X-linked genes were analysed did not show any significant associations using a Bonferroni-adjusted significance threshold of 4.5e^-5^ (Table 5). The most significant association for the combined analysis was in the chromosome X open reading frame 51B (*CXorf51B)* (p-value = =1.28e^-4^) coding for an uncharacterised protein (LOC100133053). The most significant association for males was in an RNA coding region that interacts with Piwi proteins (*DQ590189.1,* p-value = 1.7e^-3^), a subfamily of Argonaute proteins. While Piwi proteins are involved in germline stem cell maintenance and meiosis the function of the Piwi interacting RNA molecules are unknown (56). For females the most significant gene was *ARMCX1* (p-value = 6.07e^-4^) a tumour suppressor gene involved in cell proliferation and apoptosis of breast cancer cells. While this gene has not been previously implicated in TB susceptibility, *M. tuberculosis* has been shown to affect apoptosis pathways in order to evade the host immune response, suggesting that *ARMCX1* could affect TB susceptibility (57). While not significant the analysis again reveals strong sex specific effects and the sex-stratified and combined analysis gave three different results (Table 5).

**Table 5:**
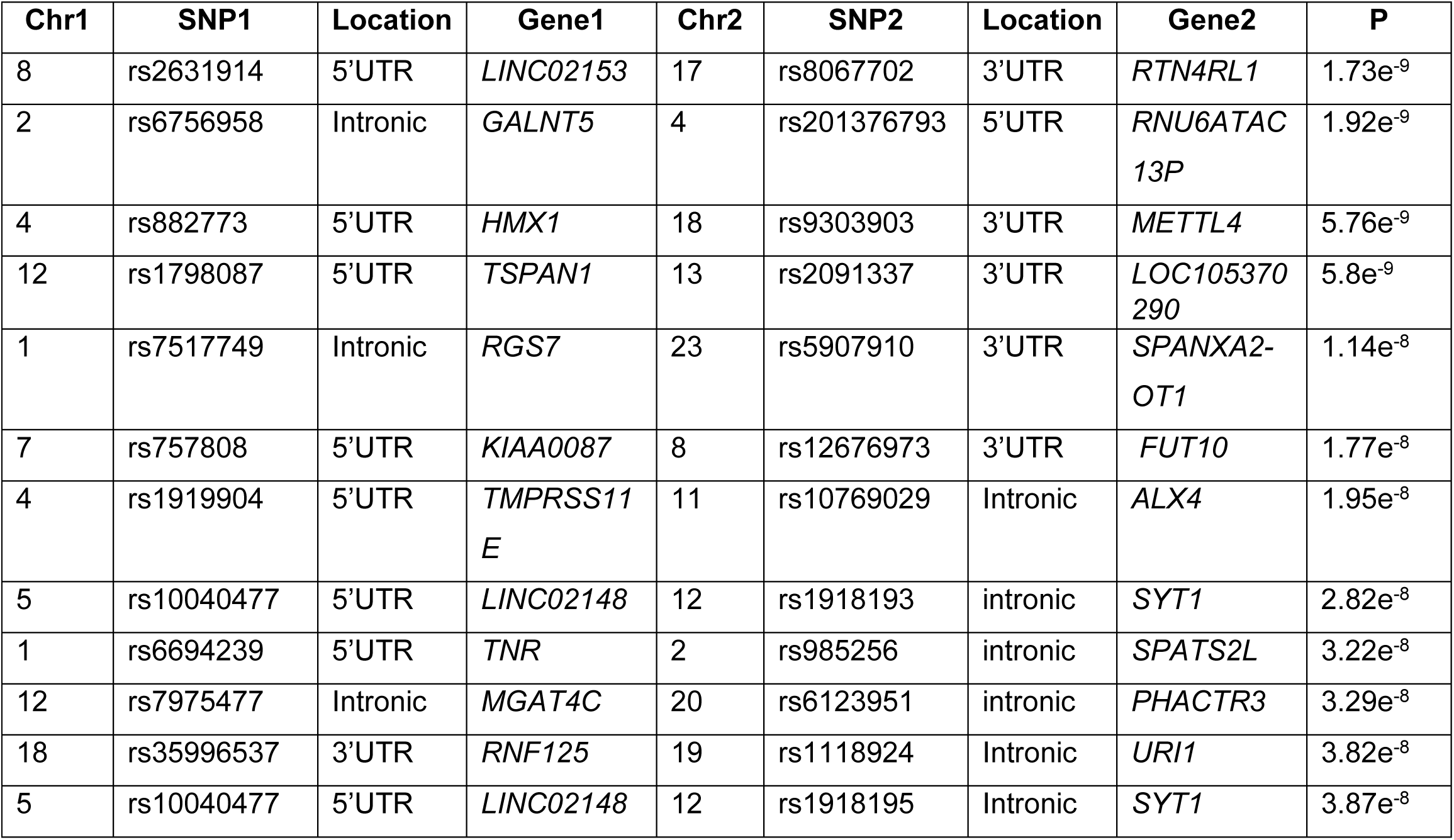
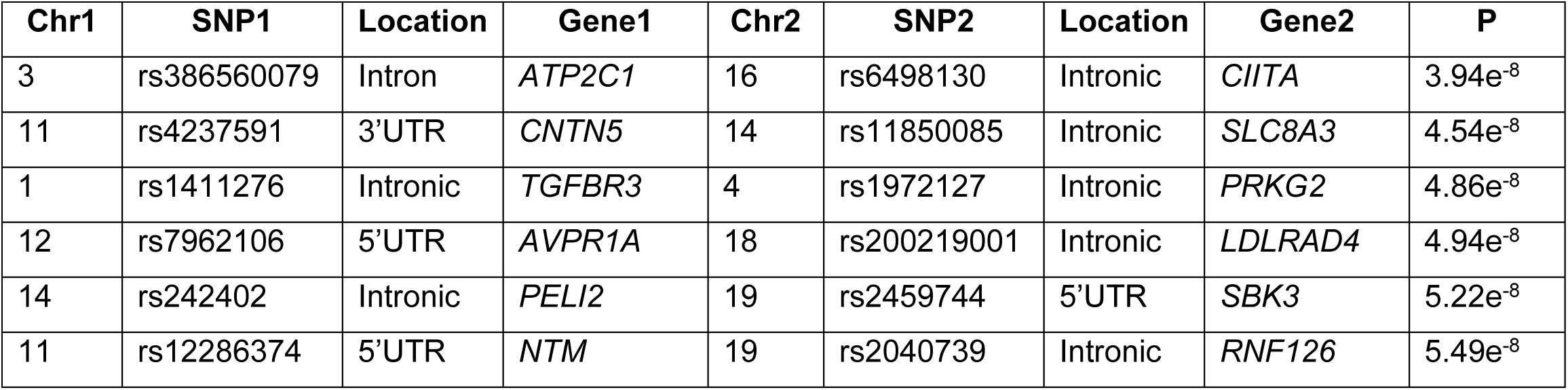
Logistic regression interaction analysis with covariate adjustment.

| Chr1 | SNP1 | Location | Gene1 | Chr2 | SNP2 | Location | Gene2 | P |
| --- | --- | --- | --- | --- | --- | --- | --- | --- |
| 8 | rs2631914 | 5'UTR | *LINC02153* | 17 | rs8067702 | 3'UTR | *RTN4RL1* | 1.73e<sup>-9</sup> |
| 2 | rs6756958 | Intronic | *GALNT5* | 4 | rs201376793 | 5'UTR | *RNU6ATAC13P* | 1.92e<sup>-9</sup> |
| 4 | rs882773 | 5'UTR | *HMX1* | 18 | rs9303903 | 3'UTR | *METTL4* | 5.76e<sup>-9</sup> |
| 12 | rs1798087 | 5'UTR | *TSPAN1* | 13 | rs2091337 | 3'UTR | *LOC105370290* | 5.8e<sup>-9</sup> |
| 1 | rs7517749 | Intronic | *RGS7* | 23 | rs5907910 | 3'UTR | *SPANXA2-OT1* | 1.14e<sup>-8</sup> |
| 7 | rs757808 | 5'UTR | *KIAA0087* | 8 | rs12676973 | 3'UTR | *FUT10* | 1.77e<sup>-8</sup> |
| 4 | rs1919904 | 5'UTR | *TMPRSS11E* | 11 | rs10769029 | Intronic | *ALX4* | 1.95e<sup>-8</sup> |
| 5 | rs10040477 | 5'UTR | *LINC02148* | 12 | rs1918193 | intronic | *SYT1* | 2.82e<sup>-8</sup> |
| 1 | rs6694239 | 5'UTR | *TNR* | 2 | rs985256 | intronic | *SPATS2L* | 3.22e<sup>-8</sup> |
| 12 | rs7975477 | Intronic | *MGAT4C* | 20 | rs6123951 | intronic | *PHACTR3* | 3.29e<sup>-8</sup> |
| 18 | rs35996537 | 3'UTR | *RNF125* | 19 | rs1118924 | Intronic | *URI1* | 3.82e<sup>-8</sup> |
| 5 | rs10040477 | 5'UTR | *LINC02148* | 12 | rs1918195 | Intronic | *SYT1* | 3.87e<sup>-8</sup> |
| 3 | rs386560079 | Intron | *ATP2C1* | 16 | rs6498130 | Intronic | *CIITA* | 3.94e<sup>-8</sup> |
| 11 | rs4237591 | 3'UTR | *CNTN5* | 14 | rs11850085 | Intronic | *SLC8A3* | 4.54e<sup>-8</sup> |
| 1 | rs1411276 | Intronic | *TGFBR3* | 4 | rs1972127 | Intronic | *PRKG2* | 4.86e<sup>-8</sup> |
| 12 | rs7962106 | 5'UTR | *AVPR1A* | 18 | rs200219001 | Intronic | *LDLRAD4* | 4.94e<sup>-8</sup> |
| 14 | rs242402 | Intronic | *PELI2* | 19 | rs2459744 | 5'UTR | *SBK3* | 5.22e<sup>-8</sup> |
| 11 | rs12286374 | 5'UTR | *NTM* | 19 | rs2040739 | Intronic | *RNF126* | 5.49e<sup>-8</sup> |

### Interaction analysis

A genome-wide interaction analysis was performed using the software Cassie. In total 1893973105 interactions were analysed and following a Bonferroni correction for the number of interactions performed the significance threshold was set to 2.6e^-11^. For the joint effects model, 18 interactions passed the significance threshold (Table S2). The most significant interaction being between rs1823897 upstream of the *ARSF* gene and rs7064174 in the *FRMPD4* gene (p-value = 7.23e^-14^), two genes for which not much information is available and it is unclear how they could be involved in TB susceptibility. The top 450 associations from the joint effects model were then retested using logistic regression and the same covariates as the SNP based association testing. No significant interactions (threshold of 2.6e^-11^) were observed in the logistic regression model (Table 6), but as Bonferroni correction is very conservative the top interactions should still be considered as they reach the significance level for SNP based GWAS.

Among the top hits in the logistic regression analysis (Table 6) some could impact TB susceptibility as they are involved in immune functions. The most significant interaction was between rs2631914 (*LINCO2153),* which is upregulated in people with major depressive disorder (58) and rs8067702 (*RTN4RL1),* previously associated with congenital heart disease, microcephaly and mild intellectual disability (59). While this interaction is not very informative in the context of TB three other interactions were identified that could impact TB susceptibility (Table 6).

The first interaction of interest is between *RNF125* gene (rs35996537) and *URI1* (rs1118924), involved in downregulation of CD^4+^/CD^38-^ T-cells and PBMCs in HIV-1 positive individuals and NF-kB/CSN2/Snail pathway, activated by TNFα respectively (60,61). Second the interaction between rs386560079 (*ATP2C1),* which is involved in regulation of intracellular Ca^2+^/Mn^2+^ concentrations through the Golgi apparatus (62) and rs6498130 (*CIITA).* Variants in the *CIITA* gene reduce the expression of *MHC class II* proteins and receptors resulting in an immune privilege phenotype (63). The final interaction of interest is between rs12286374 (*NTM),* which is mainly expressed in the brain and promotes neurite outgrowth and adhesion (64) and rs2040739 (*RNF126)* a ring type E3 ligase involved in the Protein B kinase pathway which has been previously implicated in glucose metabolism, apoptosis, cell proliferation and transcription (65). While none of these genes have previously been implicated in TB susceptibility the fact that some of them are involved in immune functions suggests a role in TB susceptibility

## Discussion

In this GWAS we investigated TB susceptibility in the admixed SAC population, with a specific focus on sex-bias and the X chromosome. A sex-stratified QC protocol was applied to the data in order to conserve inherent differences between the sexes and all statistical analysis were conducted in a sex-stratified and combined dataset in order to fully assess the impact of sex on TB susceptibility and the male sex-bias it presents with. We found no significant associations on the autosome or X chromosome for both the sex-stratified and combined SNP and gene-based association testing. A few significant interactions were identified, but the impact of these on TB susceptibility is unclear and will require further investigation to validate and functionally verify.

For the combined autosomal SNP based association testing the only potential variant of interest is rs17410035 located in the *DROSHA* gene (Table 2) which is potentially involved in miRNA biogenesis and could impact TB susceptibility if immune related regulatory miRNA is affected. For the X-linked association testing the most significant association in males was in an uninformative pseudogene, while the female and combined analysis revealed the same variant, rs768568 located in the *TBL1X* gene (Table 3). The TBL1X protein has been shown to be a co-activator of NF-kB mediated transcription of cytokine coding genes, but the mechanism of activation is unclear (53). NF-kB is a vital component of the proinflammatory signalling pathway and is involved in multiple immune pathways including TLRs (66), which have previously been shown to influence TB susceptibility (8). Based on this one could extrapolate that variants in the *TBL1X* gene could affect activation and proinflammatory signalling of NF-kB, which could have a direct effect on the immune system and thus TB susceptibility. The direction of effect for this variant was the same in males and females (Table 3), but was less significant in males probably due to loss of power when analysing haploid genotypes. For the variants identified in the sex differentiated analysis it is unclear how they could influence TB susceptibility as the most significant variant is located in a pseudogene. However, the sex differentiated test did reveal just how big the difference in effects can be between the sexes for a specific variant (Table 4). If these variants with opposite effects are not analysed in a sex-stratified way then the effects would cancel each other out and any information on sex specific effects would be lost. The X-linked gene-based association test revealed no significant associations despite having more power than the SNP based association testing. A possible reason for this could be that Bonferroni correction was used and as this is very conservative possible associations could have been missed. When looking at the most significant associations (Table 5) however it is unclear how the identified genes could be implicated in TB susceptibility.

The joint effects interaction analysis revealed several significant interactions, but as association results have been previously shown to be severely influenced by admixture (67) only the results for the logistic regression analysis will be discussed here. A few variants were identified in the logistic interaction analysis that could impact TB susceptibility (Table 6). *URI1* (rs1118924) is activated by TNFα and is involved in the NF-kB/CSN2/Snail pathway, *CIITA* (rs6498130) impacts expression of MHC class II proteins and receptors and rs35996537 (*RNF125)* and rs2040739 *(RNF126)* are both E3 ubiquitin ligase proteins which affect a multitude of cellular functions, such as apoptosis (65) and protein degradation (68). NF-kB, TNFα, MHC class II, E3 ligases, apoptosis and T-cells have all been implicated in TB susceptibility and could collectively contribute by influencing the immune response (57,68–74). As TB is a complex disease all potential influential factors need to be considered and as such the interaction analysis cannot be ignored. Shortcomings of the interaction analysis are that they are very computationally intensive and suffer from a massive multiple test correction burden. Future research should thus focus on ways to prioritise variants for interaction analysis to decrease computation time as well as have sufficient sample size to minimise multiple test correction burden.

A previous GWAS in the SAC population found a significant association with TB susceptibility in the *WT1* gene (rs2057178, OR = 0.62, p-value = 2.71e^-6^) (75). This association did not reach genome-wide significance in our study (OR = 0.75, p-value = 0.049). At the time of the GWAS by Chimusa *et al*. (75) there were few African and KhoeSan (only 6 KhoeSan) individuals in the reference data used for imputation and the accuracy of imputation in this population was not known. As the identified variant (rs2057178) was imputed into the data it should have been validated in the SAC population using an appropriate genotyping approach. Secondly although the variant reached a significance threshold for the number of variants tested it did not reach genome wide significance threshold of 5.0e^-8^ (48). Finally, the GWAS performed by Chimusa *et al.* (75) only contained 91 control individuals compared to 642 cases, which could affect the power of the study. Chimusa *et al.* (75) were unable to replicate previous associations identified in the X-linked *TLR8* gene (27). The two *TLR8* variants in our data, rs3764880 (OR = 1.73, p-value = 3.1e^-4^) and rs3761624 (OR = 1.70, p-value = 3.94e^-4^) also did not show significant associations. While the haploid genotypes in males contributes to this, a second influential factor could be admixture. Chimusa *et al.* (75) did not perform X chromosome specific admixture analysis, which could affect association testing of X-linked genes. Furthermore, only six KhoeSan reference individuals were available, which could affect the accuracy of admixture inference and severely affect the results. For our study 307 KhoeSan individuals were available, improving the admixture inference and could explain why lower stronger effects were detected for the *TLR8* variants when compared to Chimusa *et al.* (75). Candidate genes identified in previous GWAS studies were also separately analysed here, but associations did not replicate (Online supplementary material).

We did not find any significant associations with TB susceptibility, but highlight the need for sex-stratified analysis. Closer inspection of the data revealed that a large number of SNPs with opposite direction of effects for not only the X chromosome, but the autosome too. Sex specific effects has previously been reported for autosomal variants associated with pulmonary function in asthma (76). In the SAC population these opposite effects have previously been observed for X-linked variants in the *TLR8* gene (31) and the same is observed in this study. Sex-stratified analysis should therefore be included in association studies and incorporated in the study design. This can be done by keeping the male to female ratio balanced in the cases and controls. It would also be prudent to do the power calculation for the males and females separately. This will ensure sufficient power for sex-stratified analysis and could elucidate informative sex specific effects. This study was done in a 5-way admixed population. As was observed for the interaction analysis including admixture components significantly changes the association results. Furthermore it was observed that the ancestral distribution between the X chromosome and autosome are different (Figure 1), which is an indication of sex-biased admixture (43,77) and highlights the importance of including X chromosome admixture components for X-linked and sex-bias analysis. It is important to note here that the ancestral components in the SAC present with a very wide range (Figure 2) and all this variability could affect the power of association studies. It is therefore desirable to increase the sample size when analysing admixed individuals. Alternatively, a meta-analysis can be conducted, including data from all 5 ancestral populations.

**Figure 2:**
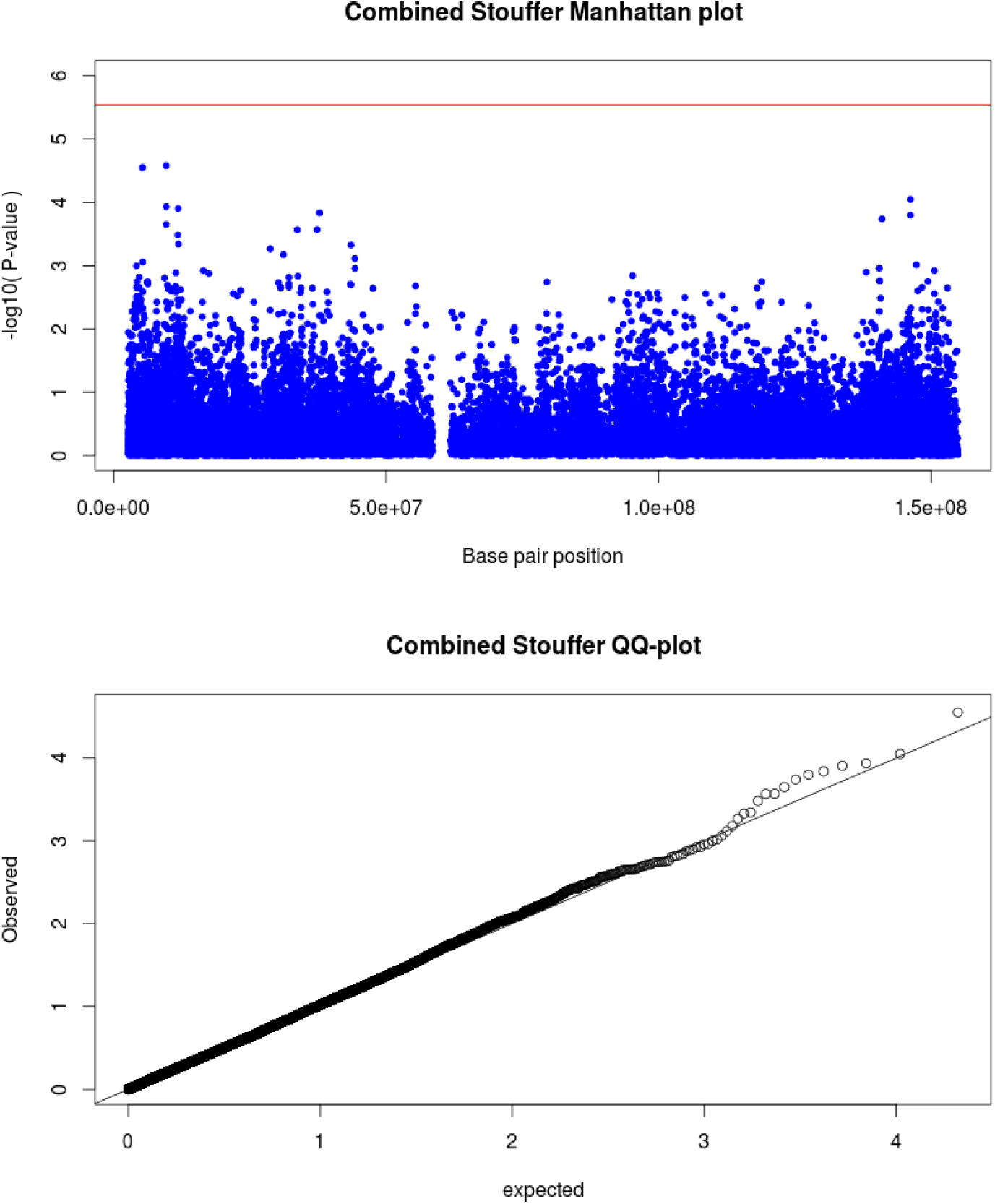
Manhattan plot (above) for X-linked associations with significance threshold indicated (red line). QQ-plot (below) shows good correlation between expected and observed p-values.

In conclusion, while no significant associations were identified this study shows the importance of conducting sex-stratified analysis. This analysis should be incorporated during the study design phase to ensure sufficient power and allow the inclusion of covariates with sex specific effects (in this case admixture components. The sex-stratified analysis revealed that the effect of certain variants can differ between males and females, not only for the X chromosome but also for the autosome. TB is a complex disease with most genetic associations that do not replicate across different populations, which complicates the elucidation of the genetic impact on disease susceptibility. By including sex-stratified analysis and identifying sex specific effects and the cause for the male bias we can adjust treatment according to sex and potentially improve treatment outcome and survival.

### List of abbreviations

95CI: 95% confidence interval
ARMCX1: Armadillo repeat containing X-linked 1
ARSF: Arylsulfatase F
ASNS: Asparagine synthetase
ATP2C1: ATPase secretory pathway Ca2+ transporting 1
C5orf64: Chromosome 5 open reading frame 64
CFAP54: Cilia and flagella associated protein 54
Chr: Chromosome
CIITA: Class II major histocompatibility complex transactivator
CXorf51B: Chromosome X open reading frame 51B
DIAPH2: Diaphanous related formin 2
DNA: Deoxyribonucleic acid
DPF3: Double PHD fingers 3
DROSHA: Drosha ribonuclease III
FRMPD4: FERM and PDZ domain containing 4
GRAMD2B: GRAM domain containing 2B
GWAS: Genome-wide association study
HIV: Human immunodeficiency virus
HWE: Hardy-Weinberg equilibrium
IL-10: Interleukin 10
LD: Linkage disequilibrium
LINC00400: Long intergenic non-protein coding RNA 400
LINC02246: Long intergenic non-protein coding RNA 2246
LINC02153: Long intergenic non-protein coding RNA 2153
MAF: Minor allele frequency
MEGA: Multi-ethnic genotyping array
MHC: Major histocompatibility complex
MIR514A1: MicroRNA 514a-1
miRNA: micro RNA
MTND6P12: MT-ND6 pseudogene 12
NCS1: neuronal calcium sensor 1
NF-kB: Nuclear Factor kappa-light-chain-enhancer of activated B cells
NTM: Neurotrimin
OR: Odds ratio
P_comb: Combined p-value using Stouffers method
P_Diff: P-value for sex-differentiation test
PBMC: Peripheral blood mononuclear cell
PCSK6: Proprotein convertase subtilisin/kexin type 6
pTB: Pulmonary Tuberculosis
RNA: Ribonucleic acid
RN7SKP120: RNA, 7SK small nuclear pseudogene 120
RNF125: Ring finger protein 125
RNF126: Ring finger protein 126
RNU6-974P: RNA, U6 small nuclear 974, pseudogene
RTN4RL1: Reticulon 4 receptor like 1
SAC: South African Coloured
SALL2: Spalt like transcription factor 2
SNP: Single nucleotide polymorphism
SRPX: Sushi repeat containing protein X-linked
TB: Tuberculosis
TBL1X: Transducin beta like 1 X-linked
TENT4A: Terminal nucleotidyltransferase 4A
TLR: Toll-like receptor
TNFα: Tumor necrosis factor alpha
TST: Tuberkulin skin test
URI1: URI1, prefoldin like chaperone
WT1: Wilms tumor 1
XWAS: X chromosome wide association study

## Acknowledgements

We would like to acknowledge and thank the study participants for their contribution and participation. This research was partially funded by the South African government through the South African Medical Research Council. The content is solely the responsibility of the authors and does not necessarily represent the official views of the South African Medical Research Council. This work was also supported by the National Research Foundation of South Africa (grant number 93460) to EH. This work was also supported by a Strategic Health Innovation Partnership grant from the South African Medical Research Council and Department of Science and Technology/South African Tuberculosis Bioinformatics Initiative (SATBBI, GW) to GT.

## Author contribution

HS, MM, CK, GT conceived the idea for this study. CG, GW, BH did the calling and QC of the raw genotyping data. HS did the analysis and wrote first draft. BH Assisted with admixture analysis. All authors contributed to writing and proofreading for approval of the final manuscript.

## Footnotes

1 http://pngu.mgh.harvard.edu/purcell/plink/

2 http://www.python.org

3 https://www.staff.ncl.ac.uk/richard.howey/cassi/using.html

